# Cannabidiol Impairs Glucagon Trafficking and Secretion Through the Lysosomal System in Murine *α*TC1-6 cells

**DOI:** 10.64898/2026.10.30.762933

**Authors:** Samuel J.P. Ugulini, Daniel B. Hardy, Savita Dhanvantari

## Abstract

Glucagon secretion by pancreatic *α* cells is regulated by nutrient, paracrine, and neural inputs, and emerging evidence suggests that the islet endocannabinoid system (ECS) may also regulate *α*-cell function. While phytocannabinoids such as cannabidiol (CBD) have been shown to interact with the ECS, and negatively regulate insulin secretion from pancreatic β cells, whether CBD alters *α* cell function in ways that disrupt glucagon trafficking and secretion remains unclear. This study investigates how acute CBD exposure affects transcriptional and secretory networks governing *α* cell function. Using the *α*TC1-6 glucagon-secreting cell line, we exposed the cells to 0.3 and 3 µM CBD for RT-qPCR analysis, 3 µM CBD for bulk RNA sequencing, and 3 µM CBD or its inactive metabolite 7-COOH-CBD for imaging and secretion experiments. Transcriptomic analysis revealed that treatment with CBD did not alter the expression of ECS receptors and enzymes. However, pathways involved in cellular excitability, peptide hormone trafficking, and secretion were significantly downregulated. Using confocal microscopy, we found that CBD exposure increased the colocalization of glucagon with LAMP1 (a lysosomal marker), and that CBD rerouted peripheral glucagon, and glucagon-containing lysosomes towards the intracellular region of the cell. In conjunction with these findings, we observed reduced glucose-stimulated glucagon secretion under high glucose conditions. Overall, our data supports a model in which acute CBD exposure downregulates processes governing cellular excitability and peptide secretion and negatively regulates glucagon trafficking and secretion by rerouting peripheral glucagon into lysosome-associated intracellular compartments in *α*TC1-6 cells.

## Introduction

Concomitant with the rise in recreational cannabis use, there is emerging research into the potential of phytocannabinoid-based therapies to treat clinical disorders. Studies have demonstrated a potential role for these drugs in altering the regulation of feeding behaviour, energy homeostasis, and metabolism[1–3]. Studies in rodent pancreatic β cells provide mechanistic evidence that these compounds directly affect islet function and hormone secretion. Endocannabinoid signalling can either enhance or suppress insulin secretion, depending on the ligand, receptor, and experimental context [4–6]. Additionally, CBD treatment was shown to reduce insulin secretion in clonal beta cells, a mechanism dependent on the endocannabinoid system (ECS) receptor G-protein– coupled receptor 55 (GPR55)[7]. Despite these divergent effects, alterations in intracellular Ca²⁺ signalling appear to be a recurring mechanism underlying cannabinoid-mediated changes in β-cell function [4–6]. While these findings obtained in β cells cannot necessarily be extrapolated to glucagon-secreting *α* cells, they provide a rationale to investigate whether CBD exposure similarly influences glucagon secretion. Although CBD can modify glucose homeostasis, the endocrine mechanisms underlying this modulation remain poorly understood. Particularly, whether CBD directly influences pancreatic *α* cell function and glucagon secretion has yet to be explored. Given that glucagon is a central regulator of hepatic glucose production [8–10], and an essential component of the counterregulatory response to hypoglycemia[11,12], examining the effects of CBD on glucagon secretion could further clarify the mechanisms through which CBD influences metabolic regulation.

Glucagon secretion from pancreatic *α* cells is tightly regulated by nutrient availability, paracrine hormones, and neurotransmitters[13], with emerging evidence suggesting that *α* cell function may also be regulated by the ECS. The ECS consists of endogenous ligands—anandamide (AEA) and 2-arachidonoylglycerol (2-AG)—which are synthesized and degraded by the enzymes N-acyl phosphatidylethanolamine phospholipase D (NAPE-PLD), diacylglycerol lipase (DAGL), fatty acid amide hydrolase (FAAH) and monoacylglycerol lipase (MAGL). The ECS also includes several receptors, such as cannabinoid receptors 1 and 2 (CNR1/CNR2), GPR55, and the ionotropic transient receptor potential vanilloid 1 (TRPV1) channel. CNR1 and CNR2 signal predominantly through G*α*i/o-dependent mechanisms that can reduce cAMP accumulation and modulate Ca²⁺ and K⁺ channel activity [14–16]. In contrast, GPR55 and TRPV1 promote membrane depolarization and intracellular Ca²⁺ signalling through indirect second-messenger pathways and direct ion-channel activation, respectively [17,18]. These pathways targeted by ECS receptors are all important regulators of islet cell excitability, vesicular trafficking and hormone secretion [13,19,20], providing plausible mechanisms through which cannabinoids such as CBD could alter *α* cell function and glucagon

Regulation of glucagon secretion does not only occur at the level of secretory granule (SG) exocytosis, but also through intracellular mechanisms that control glucagon synthesis, trafficking, and degradation[13,21–23]. Work from our group has identified a novel lysosome-dependent mechanism regulating glucagon availability within *α* cells, where SGs are targeted for degradation rather than released [22]. Using an *α* cell model, we demonstrated that peripheral glucagon colocalizes with lysosomes at the plasma membrane[22], and that treatment with the paracrine signals insulin/GABA and somatostatin promoted the retrograde redistribution of lysosomes from the cell periphery toward intracellular regions, linking lysosomal positioning with changes in glucagon trafficking and secretion[24]. Although lysosomal peptide trafficking has been shown to regulate insulin granule turnover in pancreatic β cells[25] and is recognized in several peptide hormone-secreting cell types[26], whether CBD-related signalling regulates this lysosome-dependent pathway in *α* cells remains unknown.

CBD has been reported to modulate autophagy-related, lysosome-associated, and apoptotic pathways through ECS receptor-dependent signalling, ERK1/2 and AKT activity, and changes in ion-channel function[27–29]. In one study, CBD increased autophagosome abundance, the precursor vesicle which fuses with lysosomes to facilitate degradation of engulfed material [27]. Additionally, antagonism of CNR1, CNR2, and TRPV1 attenuated this response, supporting the involvement of these receptors in CBD-induced autophagy[27]. However, these effects have been characterized primarily in neuronal and cancer cell models [27–29], and whether CBD alters lysosomal trafficking in pancreatic *α* cells is unknown. Because pathways regulating cellular excitability and vesicular trafficking are central to glucagon secretion and overlap with signalling networks affected by CBD[7,13,19,20,30], we hypothesized that CBD exposure alters the lysosomal trafficking of glucagon, thereby impairing glucagon secretion. Specifically, we examined whether CBD exposure was associated with glucagon redistribution toward lysosome-associated compartments and impaired glucagon secretion. To test this hypothesis, we used the mouse-derived *α*TC1-6 glucagon-secreting cell line and combined RT-qPCR, bulk RNA sequencing, confocal microscopy, and glucagon secretion assays to characterize CBD effects on ECS components, *α*-cell transcriptional networks, lysosomal trafficking, and glucagon secretion.

## Materials and Methods

### Cell Culture

Alpha TC1 clone 6 (*α*TC1-6) cells (a kind gift from C. Bruce Verchere, University of British Columbia, Vancouver, BC, Canada, ATCC no: CRL-2934™, RRID:CVCL_B036), are a clonal derivative of the parental *α*TC1 line, which originated from a pancreatic adenoma in transgenic mice expressing SV40 large T antigen under the control of the rat preproglucagon promoter. Cells were maintained in 25 mM glucose Dulbecco’s modified eagle medium (DMEM) with GlutaMAX™ Supplement, (cat#10569044, Gibco^TM^) supplemented with 15% horse serum (HS), 2.5% fetal bovine serum (FBS), and 100 units/ mL penicillin-streptomycin (Life Technologies) at 37°C and 5% CO_2_.

For all experiments, cells were seeded below passage 10. To determine cell viability, cells were seeded at 10,000 cells per well in 96-well plates. For qPCR, microscopy, and secretion experiments, cells were seeded at a density of 100,000 cells per well in 6-well plates, and for RNA-seq, cells were seeded at 1,000,000 cells per 10 cm dish. Cells were grown to 70-80% confluency (20% for cell viability and 50% for imaging experiments) in high 25 mM glucose medium supplemented with GlutaMAX™ and were incubated in serum free media with GlutaMAX™ supplement before and during phytocannabinoid treatment. For basal glucagon secretion, phenol-, serum-, and glucose-free DMEM media (cat#A1443001, Gibco^TM^) was supplemented with GlutaMAX™ (cat#35050061, Gibco^TM^), sodium pyruvate (cat#11360070, Gibco^TM^) and with high (25 mM) glucose, (cat#A2494001, Gibco^TM^).

### Cannabidiol Treatment

To determine an appropriate dosing regimen, cells were seeded at a density of 10,000 cells/well in 96-well plates and treated with increasing concentrations of 0.3, 1, 3, 5, 10, 15, 25, and 30 µM of cannabidiol (CBD, cat#90080, Cayman Chemicals, Ann Arbor, MI), and vehicle control (0.005% EtOH) (n=6) for 24 hours. CBD cytotoxicity was assessed by adding 10 µL/well of the WST-8 Assay Kit (cat#65475, abcam) and incubating for 24 hours. Absorbance was read at 460nm using an Eon microplate reader and Gen5 software (BioTek). Cell viability was calculated as a percentage relative to the vehicle control, which was designated 100% viable. Appropriate doses of CBD were selected based upon the half-maximal inhibitory (IC_50_) values generated from the cytotoxicity assay **(Figure 1**) and were supported by doses used in current literature[31]. For all experiments, cells were treated with either 0.3, or 3 µM of CBD, or vehicle control (0.005% EtOH, unless otherwise stated). For imaging and secretion experiments, 3 µM of 7-COOH-CBD (cat#21637, Cayman Chemicals, Ann Arbor, MI) was used as a negative control due to its low binding affinity/high inhibition constant (K_i_) to, and negligible activation of canonical ECS receptors[32,33], where 3 µM was chosen to match the dosage of CBD.

**Figure 1.**
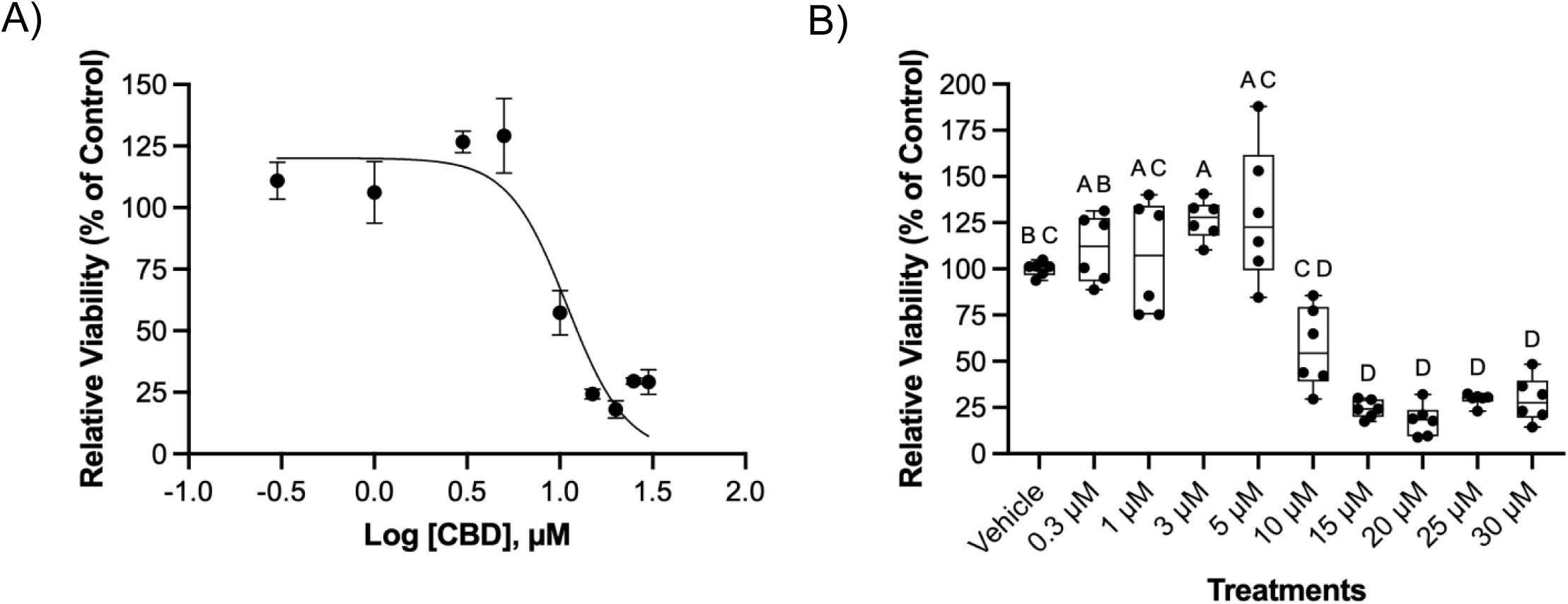
CBD reduces *α*-TC1-6 cell viability at concentrations ≥10 µM. *α*-TC1-6 cells were exposed to vehicle or CBD (0.3–30 µM) for 24 hours (n=6), followed by a 24-hour incubation with the WST-8 Assay Kit (cat#65475, abcam) to assess cell viability. A) Four-parameter logistic dose–response curve fit to the CBD-treated groups, data reported as means ± SEM. The vehicle control was excluded from curve fitting. The estimated IC_50_ was 9.742 µM (95% CI: [8.501–10.830] µM). B) Cell viability at each CBD concentration is expressed as a percentage of the vehicle control. Data points represented as means and were analyzed via one-way Welches ANOVA with Dunnet’s T3 post hoc test. Different letters indicate statistical significance between groups (p<0.05), whereas groups sharing letters are not significantly different.

### RNA extraction and RT-qPCR

For qPCR experiments, *α*TC1-6 cells were treated with 0.3 µM or 3 µM of CBD (n=8) or vehicle control (0.0005%, or 0.005% EtOH respectively (n=7) for 24 hours. Vehicle-only controls were evaluated separately to assess potential vehicle-dependent variability. Because outcomes did not differ between vehicle concentrations, vehicle controls were combined for downstream statistical comparisons (n=14). After the treatment period, cells were collected in 1 mL TRIZOL (cat#T9424, Sigma-Aldrich®), and flash-frozen at - 80^°^C. Extracts were thawed on ice, and RNA was extracted using a 1:4 ratio of chloroform (cat#C2432, Sigma-Aldrich®):TRIZOL. For each sample, 2 μg of RNA was converted to cDNA using a high-capacity cDNA reverse Transcription Kit (cat#4368814, Thermo Fisher Scientific; Applied Biosystems) following the manufacturer’s instructions using a GeneExplorer Thermal Cycler 96 (Bioer). qPCR experiments were carried out at 1:40 cDNA dilutions. mRNA expression of ECS, and other metabolic and secretory genes of interest, were determined by RT-qPCR using SensiFAST SYBR No-ROX Mix (cat#BIO-98005, Meridian Bioscience) and the CFX Opus 384 Real-Time PCR System (BioRad) using the following protocol: an initial denaturation at 95°C for 3 min followed by 43 cycles of 95°C for 15 s, 58°C for 30 s, and 72°C for 30 s. All primers are listed in **Supplementary Table 1**. Melt peaks for each primer were assessed to check for one distinct peak, and primers were tested for efficiency using a 6-fold dilution series; primers with 90-110% efficiency were chosen for gene amplification. All genes were normalized to the geometric means of actin-related protein 2 (*Actr2)* and hypoxanthine phosphoribosyltransferase 1 *(Hprt1)* housekeeping genes using the 2^−*ΔΔC*^*^T^* normalization method. These genes were chosen as their expression remained unchanged across all treatment groups.

### Bulk *α*TC1-6 cell RNA-sequencing

For RNA-seq analysis, total RNA from *α*TC1-6 cells treated with CBD (3 µM) or vehicle control for 24 hours (n=5) were assessed. Frozen *α*TC1-6 cells in TRIZOL (n=5 per treatment) were sent to Genome Quebec (Montreal, Quebec, Canada) for total RNA extraction, library preparation, and RNA-Seq. RNA quality was assessed using the RNA integrity number (RIN). All RIN scores were ≥ 7.0. Paired-end reads (25 million, 100 bp) were sequenced on the Illumina NovaSeq platform. Raw reads were pre-aligned and annotated with the UCSC mm10 mouse genome (GRCm38-mm10) using Illumina DRAGEN (software v4.3.13). Raw counts were generated using the Rsubread subpackage featureCounts in R(4.5.0). Normalization and differential expression (DE) analysis were done using the DSeq2 R package[34]. Briefly, gene-level count data were analyzed for differential expression using DESeq2. Counts were normalized using DESeq2’s default *median-of-ratios* method, and differential expression was tested using DESeq2’s Wald test. To account for multiple testing, p-values were adjusted using the Benjamini–Hochberg false discovery rate (FDR) correction. Genes with FDR < 0.05 were considered to be significant.

### Functional Enrichment Analysis

To convert ENSEMBL gene IDs to gene symbols, and to perform functional enrichment analysis on the DE genes, the clusterProfiler R package[35] with org.Mm.eg.db annotations was used. The databases included in the over-representation analysis were as follows: Gene Ontology (GO) database and its three sub-categories, biological process (BP), cellular component (CC), and molecular function (MF); and the Kyoto Encyclopedia of Genes and Genomes (KEGG) database. Enriched GO terms and KEGG pathways with adjusted p-values (q-values) < 0.05 were considered significant. Enriched terms were further validated through parallel analysis using sets of up and downregulated DE genes imported into Metascape (software v.3.5). The databases included in the over-representation analysis were as follows: GO database and its three sub-categories BP, CC, MF; the KEGG database; the Reactome (REAC) database; the Comprehensive Resource of Mammalian Protein Complexes (CORUM) database; and the WikiPathways (WP) database. Expression profiles within pathways of interest were generated as reported previously[36].

### Immunofluorescence microscopy

For imaging experiments (n=5-12), cells were seeded at a density of 100,000 cells/well onto collagen-coated glass coverslips (No. 1.5). Cells were grown to a confluency of 50%, and serum-starved for 2 hours before treatment with 3 µM CBD, 3 µM 7-COOH-CBD, or vehicle control for 24 hours. Cells were then rinsed once with warm phosphate-buffered saline (PBS), and fixed with 2% paraformaldehyde (in PBS) for 30 min and permeabilized with 0.1% Triton X-100 (in PBS) for 5 min. After a 90 min incubation in blocking buffer (2% donkey serum and 0.05% Tween 20 in PBS) at room temperature (RT), cells were incubated overnight at 4°C with the appropriate primary antibodies against glucagon (Mouse monoclonal, cat#ab10988, RRID:AB_297642, Abcam), lysosomal associated membrane protein 1 (LAMP1) (Rat monoclonal, cat#ab25245, RRID:AB_449893, Abcam), synaptotagmin 5 (SYT5), (Rabbit polyclonal, cat#PA5-44987, RRID:AB_2610517, Invitrogen™), and syntaxin1a (STX1A) (Rabbit monoclonal, cat#ab272736, RRID:AB_3741520, Abcam) diluted in blocking buffer (**Supplemental Table 2**). Cells were sequentially washed twice with PBS and incubated with the appropriate fluorophore-conjugated secondary antibodies (**Supplemental Table 2**) at RT for 90 min. Nuclear counter-staining was performed using DAPI (1:1000 in PBS), and coverslips were mounted on SuperFrost Plus glass slides (cat#1255015, Fisher Scientific, Toronto) with ProLong™ Glass Antifade Mountant (cat#P36980, Invitrogen™), sealed with nail polish, and stored at 4°C until analysis.

### Confocal microscopy

Imaging was performed on the Nikon A1R confocal microscope (Canada), equipped with NIS-Elements AR 5.41.01 software, using 60X oil-immersion objective with Galvano laser, at 4024 x 4024, pinhole size (1), average (4), and dwell time (1). The microscope was equipped with four lasers, including DAPI (405 nm), FITC (490 nm-519 nm), TRITC (561-594 nm), and Cy5 (633-647 nm). Images were processed using the AI algorithm (denoise.ai) to remove noise from the images, 2D deconvolution for maximum contrast and brightness, and background was subtracted in ImageJ FIJI (v2.16.0/1.54p/Java 21.0.7 (64 bit)) using a rolling ball radius of 50 pixels. Three to four images per biological replicate were captured, and image analysis was conducted on 1-5 cells per image (4-20/replicate), with one microscope slide considered as one biological replicate.

### Cellular region definition

Cellular regions within *α*TC1-6 cells were defined using the fluorescence intensity of different known cellular region markers. Specifically, DAPI and STX1A fluorescence were used to define the nuclear and the cell periphery regions, respectively. Whole-cell boundaries were determined in QuPath (v0.7.0-arm64) using the wand and brush tools to manually outline Stx1a membrane staining to generate a cell mask for 1-4 cells/image. The peripheral region of interest (ROI) was defined as a 600 nm inward band from the cell boundary[37,38], and the intracellular ROI was generated by subtracting the peripheral ROI from the original whole-cell mask. The nucleus was defined using a DAPI-based ROI, and the nuclear area was subtracted from both the peripheral and intracellular ROIs to remove overlap between regions. Mean fluorescent intensity from each ROI was normalized to the area of the peripheral and intracellular masks, generating average density measurements for each subcellular compartment.

### Co-localization of proteins and re-distribution analysis

Images were analysed for colocalization by Image J-FIJI Just Another Colocalization plugin (JACoP, v2.1.4) with signal overlap reported using Manders Overlap (MOV) Coefficient. M1 represents the fraction of channel 1 signal overlapping with channel 2, while M2 represents the fraction of channel 2 signal overlapping with channel 1. For redistribution analysis, images from individual channels were thresholded and processed using the “Image Calculator” function on ImageJ FIJI under the “AND” operation so that only the pixels containing two wavelengths would be captured with adjusted intensity. Plot profiles of the co-localized pixels were generated, and the redistribution of the co-localized signals was quantified and categorized to the “intracellular region” or the “cell periphery”, as described above[24].

### Glucagon Secretion

For glucagon secretion experiments (n=6), *α*TC1-6 cells were treated with CBD (3 µM), 7-COOH-CBD (3 µM), or vehicle control (0.005% EtOH) for 20 hours. To assess regulated secretion, cells were incubated for 2 h to measure basal secretion. Basal media was collected into tubes containing protease inhibitor (cat#A32955, Pierce^TM^) and stored at –80°C until analysis. Cells were rinsed twice with cold PBS, scraped in 750 µL Radioimmunoprecipitation Assay (RIPA) buffer (cat#89900, Pierce^TM^) containing protease inhibitor (cat#A32955, Pierce^TM^), centrifuged at 14,000 x g for 15 minutes, and stored at –80°C until analysis. Total protein concentration was determined using the Bicinchoninic Acid Assay (BCA) (cat#23225, Pierce^TM^) kit. Glucagon concentration in the media was determined using the human glucagon Enzyme-Linked Immunosorbent Assay (ELISA) kit (cat#EHGCG, Invitrogen™) and normalized to total protein concentration.

## Statistics and Data Analysis

All statistical analyses were performed using GraphPad (version 11.0.0), with P < 0.05 as the significance threshold. We determined IC_50_ from cell viability data using four-parameter nonlinear regression, with the bottom constrained to 0 and the top left unconstrained. qPCR data were first normalized using the 2^ΔΔct^ method, and groups were compared via a one-way Brown-Forsythe and Welch’s ANOVA with Dunnett’s T3 post hoc test to determine significance between groups. To determine glucagon secretion values, we performed a sigmoidal four-parameter nonlinear regression with no constraints and normalized secretion data to their respective total protein concentrations. For microscopy and secretion analysis, data were analyzed using a one-way Brown-Forsythe and Welch’s ANOVA with Dunnett’s T3 post hoc test.

## Results

### Higher Concentrations of CBD Significantly Reduce Cell Viability in *α*TC1-6 Cells

To determine non-cytotoxic CBD concentrations for subsequent experiments, *α*TC1-6 cells were exposed to 0.3 to 30 µM CBD for 24 hours. CBD significantly reduced cell viability in a concentration-dependent manner with an IC_50_ = 9.742 (95% CI: 8.501 to 10.830; **Figure 1A**). At 10 µM, CBD reduced cell viability to 45% of vehicle control (p=0.0484), while viability was below 10% following exposure to 15-30 µM CBD (p<0.0001; **Figure 1B**).

### CBD Exposure Does Not Affect Transcript Expression of ECS Receptors or their Corresponding Synthesizing and Degrading Enzymes

Provided that cell viability was sustained at CBD concentrations below 10 µM and significantly reduced at and above 10 µM, we chose 0.3 and 3 µM as our two doses to characterize concentration-dependent changes in mRNA transcript expression. We first determined the effects of CBD (0.3 and 3 µM) on the steady-state mRNA levels of the metabolic genes glucagon (*Gcg*), glucose transporter 1 (*Glut1*), lysosome-associated membrane protein 2 (*Lamp2*), stathmin-2 (*Stmn2*), and mucolipin 1 (*Mcoln1*). We also determined transcriptional changes of ECS receptors and ligand metabolic enzymes. Treatment with either 0.3 or 3 µM CBD did not affect mRNA transcript expression of *Gcg*, *Glut1* or *Mcoln1*. However, treatment with 0.3 µM CBD significantly increased mRNA expression of the degradative lysosomal marker *Lamp2* (p=0.015) and Stathmin-2 (*Stmn2*, p=0.027), which regulates the lysosomal transport of glucagon (**Figure 2)**. In contrast, 3 µM CBD did not affect *Lamp2* and significantly reduced Stmn2 expression (p=0.002) (**Figure 2)**.

**Figure 2:**
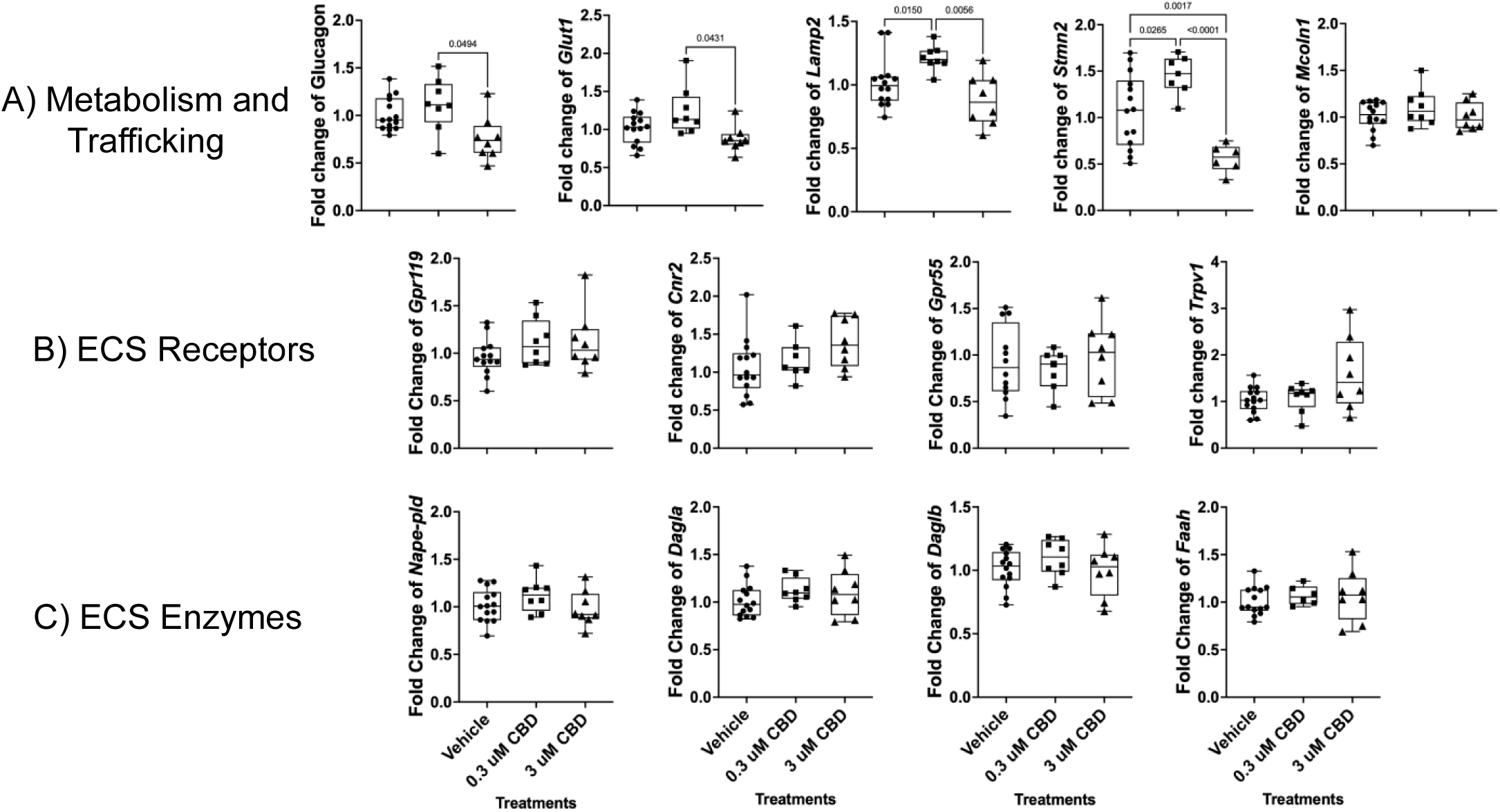
Concentration-dependent effects of CBD on gene expression in *α*-TC1-6 cells. Fold change of mRNA expression in A) genes important in alpha cell metabolism and glucagon trafficking, B) ECS receptors, and C) ECS enzymes after treatment of *α*-TC1-6 cells to vehicle (n=13-14), and 0.3 or 3 µM CBD (n=7-8) for 24 hours. All genes were normalized to the geometric means of ACTR2 and HPRT1 housekeeping genes using the 2-ΔΔCT normalization method. Data points represented as means, one-way Welches ANOVA with Dunnet’s T3 post hoc test, *p<0.05, **p<0.01, ****p<0.0001.

We also investigated the effects of CBD on mRNA expression of the ECS receptors, G-protein coupled receptor 119 (*Gpr119*), G-protein coupled receptor 55 (*Gpr55*), cannabinoid receptor 2, (*Cnr2*), transient receptor potential vanilloid 1 (*Trpv1*); and the ECS enzymes, N-acyl phosphatidylethanolamine phospholipase D (*Napepld*), fatty acid amine hydrolase (*Faah*) and DAG lipase (*Dagl*) (**Figure 2)**. There were no significant changes in the steady-state mRNA levels in any ECS receptor or enzyme upon treatment with either 0.3 mM or 3 µM CBD.

### Bulk RNA-Seq analysis of CBD-Treated *α*TC1-6 cells

Given the alternations in major *α* cell metabolic genes revealed by qPCR, we wished to investigate larger changes in *α* cell transcriptional programs altered by CBD. We conducted RNAseq analysis using cells treated with 3 µM CBD, as this dose was found to be non-toxic (**Figure 1A**), elicited strong effects on *Stmn2*, a negative regulator of glucagon secretion (**Figure 2**) and falls within the commonly used range of in vitro doses (0.1-10 µM) cited in the literature[39]. When filtering for genes with an FDR ≥ 0.05, there were 288 significantly downregulated and 174 significantly upregulated genes. An additional log2 fold change cutoff of 0.5 revealed a smaller population of 35 downregulated and 7 upregulated genes in the CBD-treated cells (**Figure 3A and B**).

**Figure 3:**
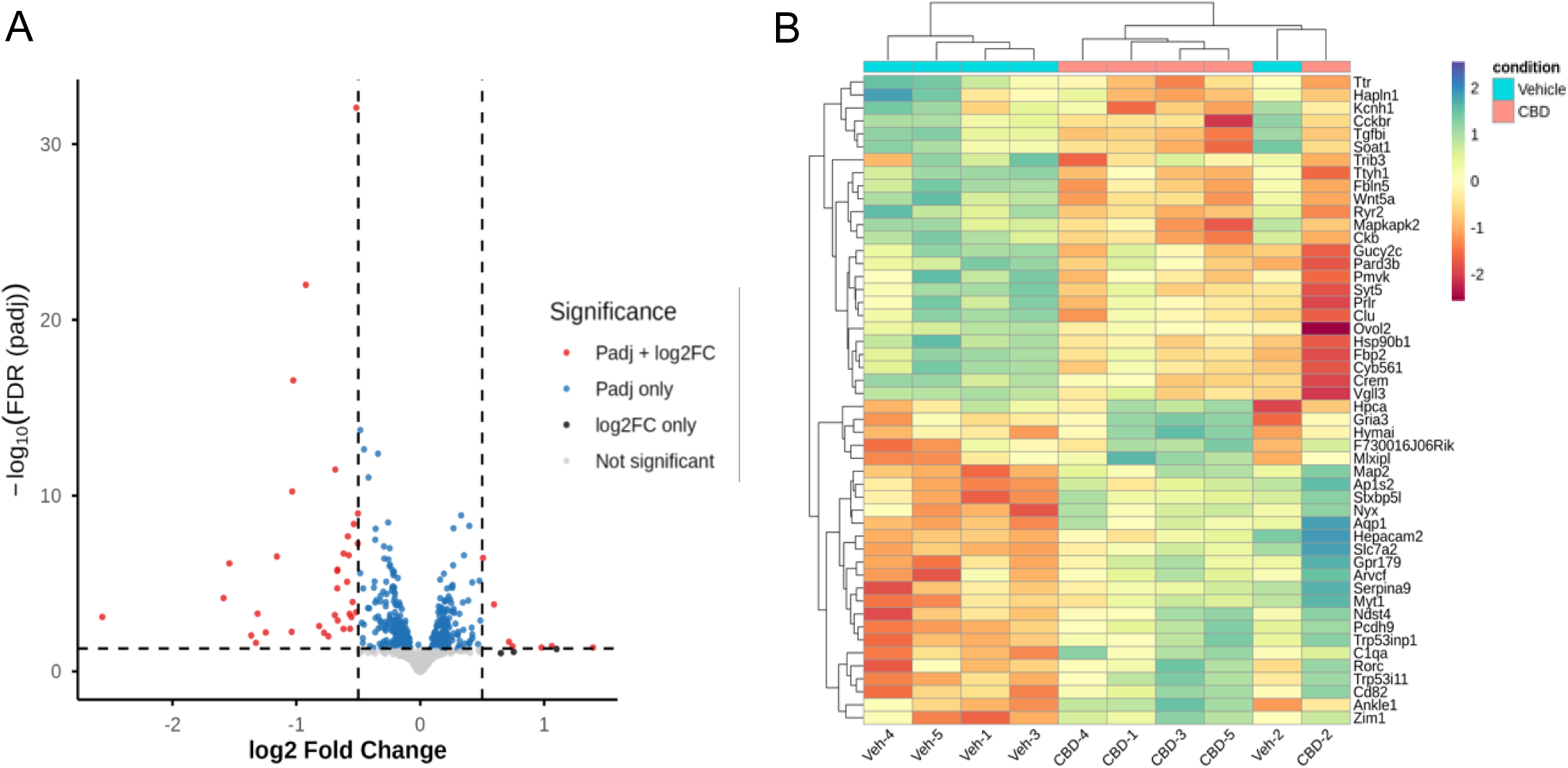
Transcriptomic changes in CBD-exposed *α*-TC1-6 cells via RNASeq. Volcano Plot A) and Heatmap B) of DE genes between vehicle- and 3 µM CBD-exposed *α*-TC1-6 Cells. RNA-Seq was performed on *α*-TC1-6 cells from vehicle-exposed (n=5) and CBD-exposed (n=5) cohorts using the Illumina NovaSeq PE100 platform with 25M reads against the mm10 genome. Differential expression analysis was conducted using Dseq2 in R. A) the log2 fold change was plotted against the log10 of the FDR for all genes in the DE comparison. Using an FDR cutoff of 0.05 there were 288 downregulated and 174 upregulated genes. When using both the FDR cutoff and a log2 fold change cutoff of 0.5 there were 35 downregulated and 7 upregulated genes. B) Heatmap was created using the log2 normalized CPM values for all DE genes (rows) and samples (columns). Values were clustered using Ward’s clustering algorithm.

### CBD Downregulates Pathways Involved in *α* Cell Excitation and Glucagon Trafficking and Secretion

Functional enrichment analysis found 853 significantly enriched terms across KEGG and GO databases. Overall, there were 308 enriched KEGG terms and 545 GO terms, encompassing 466 GO:BP terms, 36 GO:CC terms, and 43 GO:MF terms, using a p and q cutoff of 0.05 to define significance. Among the enriched terms from GO:BP, GO:CC, and GO:MF enriched terms, we observed common terms related to glucagon trafficking and secretion such as: ‘peptide hormone secretion’ (GO:BP), ‘dense core granule (GO:CC), and ‘monatomic ion channel activity’ (GO:MF) (**Figure 4**).

**Figure 4:**
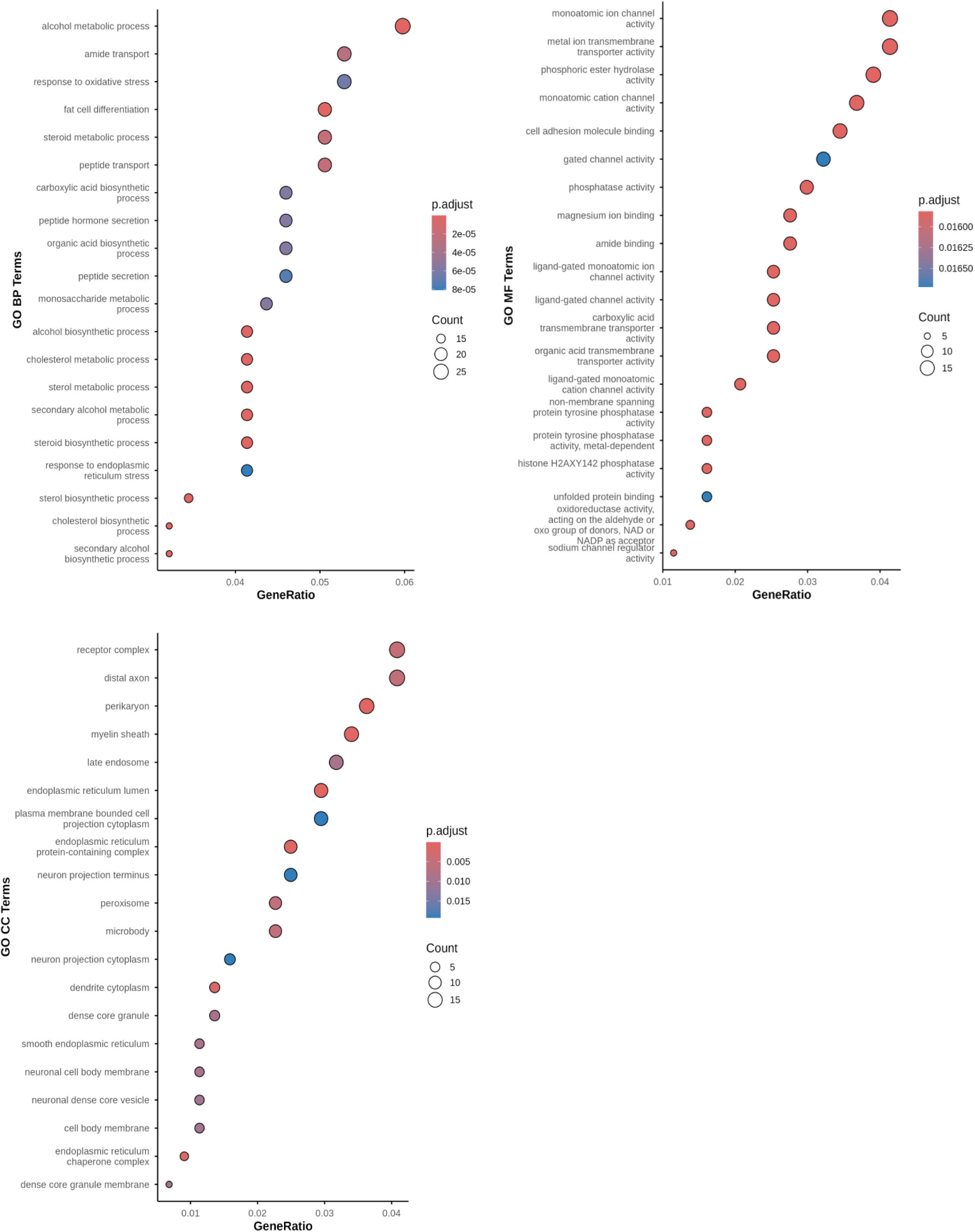
Functional enrichment analysis of CBD-treated *α*-TC1-6 cells. Dot Plots of The Top Enriched A) GO:BP, B) GO:MF, and C) GO:CC Terms. Functional enrichment analysis was performed on significantly differentially expressed genes (DESeq2, *FDR* < 0.05) using the *clusterProfiler* package (Yu et al., 2012, *OMICS*) with *org.Mm.eg.db* for mouse annotation. GeneRatio represents the proportion of DE genes that map to each GO term. Only the top 20 enriched terms are shown. Dots are coloured by adjusted pvalue (FDR) and are sized based upon the number of enriched genes in each GO term.

Across terms related to peptide hormone secretion enriched in KEGG and GO databases, we identified 14 unique genes. Five genes encoding proteins involved in ion channel activity, vesicle priming, and regulated exocytosis were upregulated (*Cacna2d1, Cadps, Kcnj6, Stxbp5l, and Adcyap1r1*), and 9 genes involved in glucagon production, ion channel activity, Ca²⁺-dependent signalling, and vesicle exocytosis were downregulated (*Kcnk1, Adcy2, Cplx1, Syt5, Kcnip4, Gcg, Plcb1, Ryr2, and Plcb1*) (**Figure 5**). Two of these genes, *synaptotagmin5* (*Syt5)–*a synaptic vesicle marker*–*and the *ryanodine receptor 2* (*Ryr2*)–involved in Ca^2+^ release from the endoplasmic reticulum, were also among the top 25 downregulated genes when accounting for significance and log2 fold change (**Figure 2B**).

**Figure 5:**
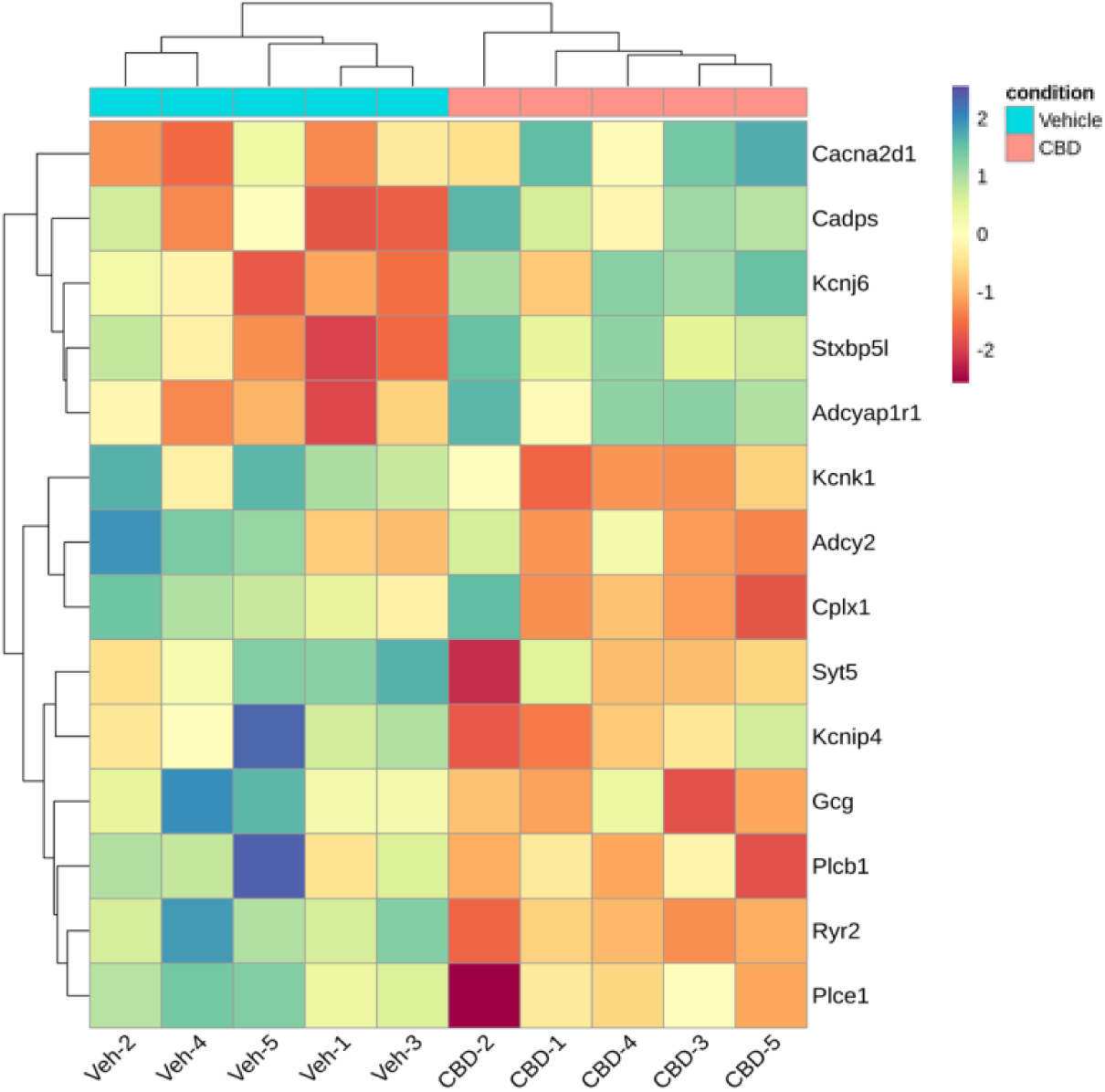
Heatmap of DE genes involved in cellular excitation and peptide hormone secretion. Functional enrichment analysis was performed on significantly differentially expressed genes (DESeq2, *FDR* < 0.05) using the *clusterProfiler* package (Yu et al., 2012, *OMICS*) with *org.Mm.eg.db* for mouse annotation. Across the GO:BP; GO:CC; GO:MF; 14 unique genes were found. The heatmap was created using the log2 normalized CPM values for all genes (rows) and samples (columns). Values were centered and scaled across rows for comparison, and clustered using Ward’s clustering algorithm.

### CBD Promotes the Intracellular Redistribution of Peripheral Glucagon

Given that (i) RNA sequencing revealed that CBD significantly downregulates SYT5, and (ii) SYT5 may be expressed on secretory granule membranes in murine *α* cells and functions in Ca2+-dependent exocytosis[40], we sought to further elucidate the functional consequences of CBD exposure on glucagon trafficking using SYT5 as a secretory granule (SG) marker. After antibody optimization, we observed a non-specific staining pattern that was uncharacteristic of our assumption that SYT5 is associated with glucagon-containing secretory vesicles. To address this, we stained both mouse and rat pancreatic islets to determine whether the current literature might be incorrect regarding the cell-type specificity of SYT5. Qualitative analysis showed that SYT5 did not localize to *α* cells in either mouse or rat islets and was instead specifically localized to β cells, with the SYT5 staining pattern closely mimicking insulin (**Supplemental Figure 1**). Based on this finding, we excluded our SYT5 staining data and proceeded without a secretory granule-specific marker. In the absence of SYT5, we investigated whether CBD affected the subcellular distribution of glucagon and lysosomes using confocal immunofluorescence microscopy (**Figure 6A**).

**Figure 6:**
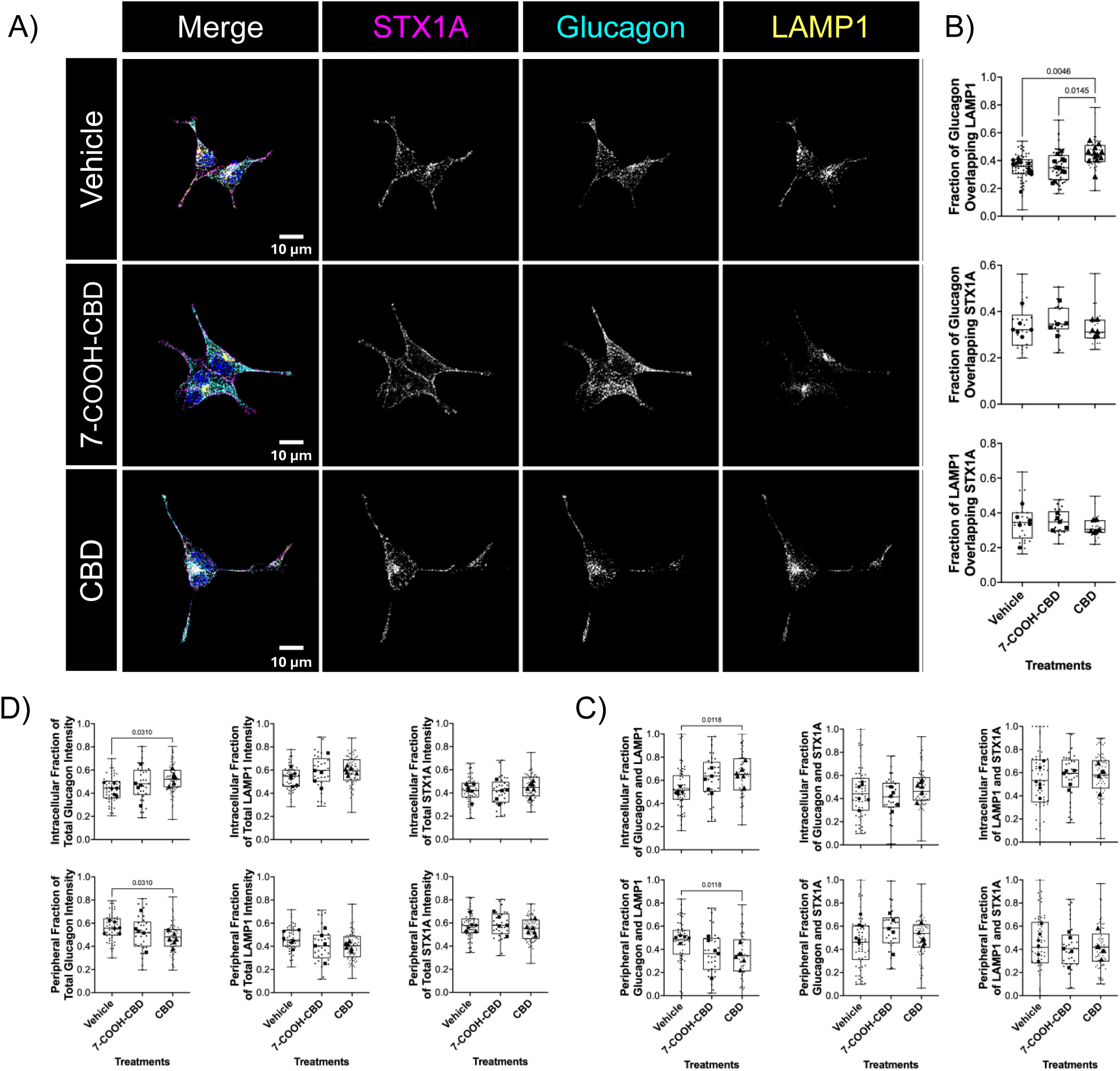
CBD increases the intracellular distribution of glucagon in lysosomes in *α*-TC1-6 cells. A) Confocal images showing the localization of glucagon, LAMP1^+^ lysosomes and with STX1A at the plasma membrane in *α*-TC1-6 cells after treatment with vehicle control, 3 µM CBD or 7-COOH-CBD for 24 hours. B) Graphs showing fractional changes in the intracellular and peripheral fractions of glucagon, LAMP1, and STX1A fluorescent intensity. D) Graphs showing fractional changes in the subcellular localization of the colocalized signal of glucagon and LAMP1, glucagon and STX1A, and LAMP1 with STX1A. D) Changes in Manders Overlap (MOV) Coefficient between glucagon and STX1A, and LAMP1 and STX1A (M1). Each large circle, square, and triangle is representative of the average fluorescent intensity or MOV coefficient of one biological replicate (n=5-6), and each small circle, square, and triangle is representative of an MOV coefficient from an individual image (3-4 per replicate), or the fluorescent intensity value from an individual cell (5-16 per replicate). Data analyzed via one-way Brown-Forsythe and Welch ANOVA with Dunnet’s T3 post hoc test. Images were processed using the Nikon NIS-Elements AI algorithm (denoise.ai) to remove noise from the image, 2D deconvolution for maximum contrast and brightness, and background intensity was subtracted in ImageJ FIJI using a rolling ball radius of 50 pixels.

Using syntaxin1a (STX1A), a soluble NSF attachment protein receptor (SNARE) protein involved in vesicle docking and fusion at the plasma membrane, as a marker of the cell periphery, we quantified changes in the subcellular localization of glucagon and LAMP1 (**Figure 6A**). Exposure to 3 µM CBD significantly increased glucagon and LAMP1 colocalization relative to both vehicle and the CBD metabolite, 7-COOH-CBD, as assessed by M1 Manders Overlap (MOV) coefficients (CBD vs vehicle, p=0.005; CBD vs 7-COOH-CBD, p<0.015, **Figure 6B**). We found that CBD treatment did not affect the distribution of glucagon or LAMP1 associated with the plasma membrane (colocalized signal of glucagon or LAMP1 with STX1A) (**Figure 6B**). All M1 coefficient and significance data can be found in Supplemental Table 2. For all comparisons, MOV M2 values were not significant across treatment groups (**Supplemental Table 3**). Provided that CBD increased the colocalization of glucagon in LAMP1^+^ lysosomes, we next examined if CBD affected the subcellular distribution of glucagon and lysosomes. We found that CBD treatment significantly increased the intracellular fraction and decreased the peripheral fractions of glucagon (p=0.031) and glucagon in LAMP1^+^ lysosomes (p=0.012; **Figure 6C/D**). Moreover, CBD treatment did not alter the subcellular distribution of either STX1A or LAMP1, nor did it affect the intracellular or peripheral fractions of glucagon and STX1A colocalization, or the LAMP1 and STX1A colocalized signal (**Figure 6C/D**). Although 7-COOH-CBD did not significantly affect the subcellular distribution of glucagon, LAMP1, or STX1A relative to vehicle, the 7-COOH-CBD group was also not significantly different from the CBD-treated group (**Figure 6C/D**).

### CBD Secretion Data

Building on these findings, we next examined whether CBD-induced transcriptional and trafficking changes translated into functional alterations in glucagon secretion. *α*TC1-6 cells were pretreated with CBD or 7-COOH-CBD for 20 hours, and exposure was maintained throughout the basal secretion assays. Interestingly, both 7-COOH-CBD and CBD significantly decreased basal glucagon secretion (p<0.0001 and p<0.001, respectively; **Figure 7**).

**Figure 7:**
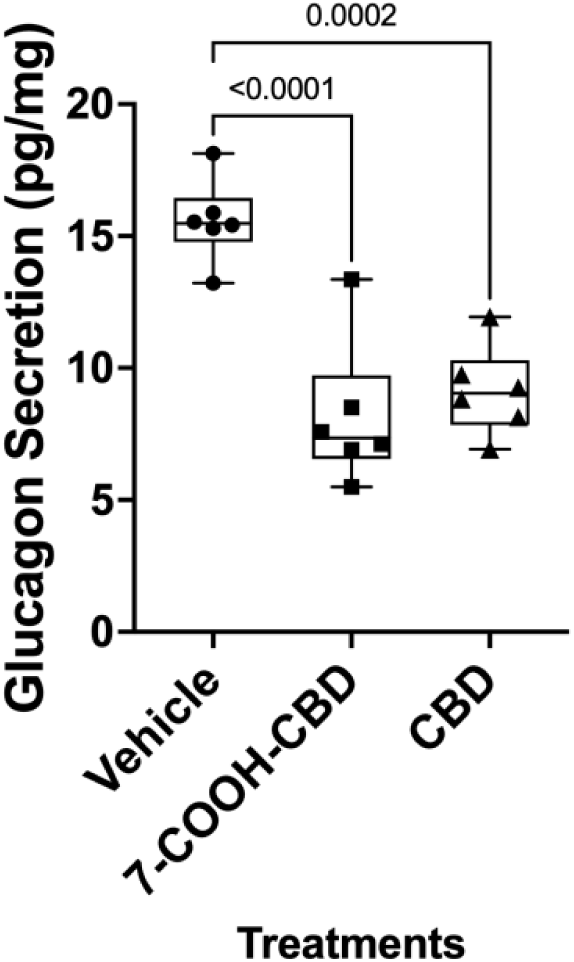
CBD reduces basal glucagon secretion in α-TC1-6 cells. Changes in basal glucagon secretion in *α*-TC1-6 cells treated with vehicle control, and 3 µM CBD or 7-COOH-CBD for 20 h. For basal secretion, cells were incubated for an additional 2 h in treatment media (n=6). Glucagon secretion was quantified by ELISA, normalized to total protein, and presented as means ± SD. Data was analyzed via one-way ANOVA with Tukey’s post hoc test.

## Discussion

While much is known regarding the mechanisms regulating secretory granule fusion and exocytosis at the plasma membrane in *α* cells[13,41,42], the contribution of intracellular transport mechanisms to glucagon trafficking and secretion has only recently begun to emerge. Previous work from our group has implicated the lysosomal system as an important regulator of glucagon trafficking [22,24,43] and demonstrated that paracrine hormones may suppress glucagon secretion through retrograde transport of glucagon-containing lysosomes[24]. Given that CBD exposure is associated with modulation of lysosome-dependent pathways via ECS signalling[27–29,44] , we investigated whether CBD could alter intracellular trafficking and secretory dynamics in *α* cells. In this study, we used transcriptomic, microscopy and functional approaches to characterize how acute CBD exposure alters glucagon trafficking and secretion dynamics through the lysosomal system.

To our knowledge, CBD-induced cytotoxicity has not been directly investigated in pancreatic islets or immortalized pancreatic cell lines such as *α*TC1-6. Nevertheless, our findings are consistent with evidence from other cell models showing that micromolar CBD can reduce cellular viability in a concentration- and time-dependent manner. CBD was found to significantly reduce cell viability to under 50% at 5 µM and above in HaCat, HeLa, MDA-MB-231 and CaCo-2 cell lines. Similarly, CBD produced 24-hour IC_50_ values of 12.3 and 16.1 µM in the ovarian carcinoma cell lines A2780 and cisplatin-resistant A2780/CP70, respectively[45]. Although these cancer-derived models are not directly comparable to pancreatic *α* cells, the findings collectively show that micromolar CBD can impair cellular viability, with sensitivity varying by cell type, exposure duration, and dose. Based on our results and those reported in the literature, we then investigated whether CBD at non-cytotoxic concentrations exerts dose-dependent effects on genes important in *α* cell metabolism and glucagon trafficking.

We first analyzed the expression of 5 genes previously identified by our lab as important to glucagon trafficking and secretion—*Mcoln1*, Glucagon, *Glut1*, *Lamp2*, and *Stmn2*. The dose-dependent effects of 0.3 µM and 3 µM CBD observed on the expression of *Lamp2* and *Stmn2* align with the positively correlated relationship between the lysosomal-associated protein LAMP2 and the microtubule-binding protein STMN2 that we have previously demonstrated in vitro (e.g. *α*TC1-6 cells) and in vivo (i.e. islets from a mouse model of T1D[22]. This paper demonstrated that in a mouse model of diabetes where STMN2 expression is significantly reduced, the colocalization of glucagon and LAMP2 is sharply decreased, suggesting a positively correlated relationship between STMN2 and LAMP2 expression. Additionally, we have previously characterized STMN2 as a negative regulator of glucagon secretion through retrograde lysosomal transport; however, in response to 3 µM CBD, *Stmn2* expression was significantly reduced, suggesting a STMN2-independent mechanism may underlie CBD-induced changes in glucagon trafficking and secretion. To further investigate changes in lysosomal biogenesis, we examined the expression of MCOLN1. This gene encodes a Ca^2+^ channel critical for the activation of lysosomal biogenesis, as it promotes Ca^2+^ availability for the activation of calcineurin, a phosphatase responsible for the activation of transcription factors (TF) TFE3/B, the master regulators of lysosomal biogenesis. A reduction in MCOLN1 would signal suppression of this pathway and subsequently point towards a reduction in lysosomal biogenesis. Additionally, we investigated changes in GLUT1 expression—the primary glucose transporter in *α* cells—as previous work from collaborators found that prenatal CBD exposure reduced GLUT1 expression in the placenta of CBD-exposed dams[46]. Upon analysis, both MCOLN1 and GLUT1 were found to be unchanged at both CBD concentrations. Provided the observed changes in metabolic and trafficking genes we next sought to characterize ECS expression in these *α*TC1-6 cells.

In addition, we found that CBD exposure, at both concentrations, did not alter ECS receptor or enzyme expression; however, there was a slight upward trend in CNR2 and GPR55 expression, a trend one would expect given CBD’s partial activity and antagonism of CNR2[47] and GPR55[48,49], respectively. Nevertheless, even though ECS transcript expression was unchanged, their detection and specific expression patterns offer novel insight into how the murine ECS functions. Our results show one of the first comprehensive transcriptomic characterizations of the ECS in murine *α* cells. However, the literature lacks consensus on cell-specific expression of ECS receptors and enzymes. Studies looking at adult mouse islets are inconsistent in their findings of ECS expression; Starowicz et al. characterized CNR1, DAGL*α* and NAPE-PLD expression mainly in *α* cells, with MAGL and FAAH being primarily localized to β cells. Kim et al. instead reported CNR1 in both *α* and β cells, CNR2 in δ cells, with NAPE-PLD, DAGL*α*, and FAAH mainly in β cells, and MAGL in both *α* and β cells. Interestingly, other studies localized CNR1 to only δ cells, whereas Malenczyk et al. found CNR1 predominantly in β cells, TRPV1 and DAGL*α* in *α* and β cells, and MAGL mainly in *α* cells. Given the inconsistent literature on ECS expression, our qPCR analysis provides a sensitive characterization of ECS transcripts in *α* cells. While we did not report CNR1 expression in *α*TC1-6 cells, this was a function of repeated attempts at finding a specific primer for CNR1 amplification, which we were unable to demonstrate, suggesting our lack of success may be attributable to a lack of expression in this cell line. This approach also addresses limitations of G-protein-coupled receptor (GPCR) immunodetection, for which antibodies require validation with appropriate positive and negative controls due to the conserved nature of GPCR proteins[50].

Because we observed transcriptional changes in genes shown to regulate glucagon intracellular trafficking and lysosomes, we assessed whether acute CBD exposure altered broader transcriptomic programs related to these processes. Across the GO:MF and GO:CC databases, genes involved in secretory vesicle exocytosis and cellular excitability were significantly downregulated. Of particular interest were glucagon, ryanodine receptor 2 (*Ryr2*), phospholipase C beta 1 and epsilon 1 (*Plcb1/Plce1*), complexin-1 (*Cplx1),* and synaptotagmin 5 (*Syt5*). RYR2 is a Ca^2+^-sensitive Ca^2+^ channel localized to the endoplasmic reticulum (ER) in pancreatic islets such as *α* and β cells[51]. RYR2 contributes to Ca^2+^-induced Ca^2+^-release upon the influx of extracellular Ca^2+^, thereby amplifying the Ca^2+^ signals needed for membrane depolarization and, thereby, glucagon secretion[51]. In addition to *Ryr2*, CBD exposure was associated with downregulation of K+ channel subunits and the Ca^2+^-dependent signalling genes *Plcb1* and *Plce1*. Together, these changes suggest that CBD may reduce cellular excitability and, consequently, hormone secretion. *Plcb1* and *Plce1* encode subunits of phospholipase C (PLC), which catalyzes the conversion of phosphatidylinositol-4,5-bisphosphate (PIP_2_) into two secondary messengers: inositol 1,4,5-trisphosphate (IP_3_) and diacylglycerol (DAG). IP_3_ then binds to ligand-gated Ca^2+^ channels (IP_3_ receptors) on the ER membrane, triggering Ca^2+^ release and contributing to glucagon secretion. Of particular interest, the ECS receptor GPR55 is known to couple to Gα_q_ and may stimulate ER Ca^2+^ release through PLC-mediated IP_3_ production. GPR55 has been shown to stimulate insulin secretion in clonal β cells[52], and reductions in these genes may be associated with CBD’s antagonism of GPR55 activity. In addition to genes involved with cellular excitability, genes involved with vesicular trafficking, such as *Cplx1* and *Syt5*, were also found to be significantly enriched/downregulated. Both CPLX1 and SYT5 are proteins known to regulate vesicle exocytosis at the plasma membrane through their interaction with SNARE proteins, promoting Ca^2+^-dependent granule release[53–56]. In contrast to CPX1, which is bound to the secretory cell membrane[53,57], SYT5 is reported to be specifically expressed on the membranes of secretory, dense-core vesicles in murine *α* cells[40].

Currently, very little is known about the function of *Syt5*, and existing data reach contradictory conclusions with respect to cellular localization in pancreatic islets. While there is evidence that SYT5 functions as a Ca^2+^-sensitive regulator of vesicle exocytosis[40,56], the nature of its cell-type specificity is debatable. Mechanistically, SYT5 binds to the syntaxin family of proteins via its C-terminal domain in a Ca^2+^-dependent manner, supporting its role in mediating membrane fusion of dense-core vesicles in neuro- and non-neural-endocrine cells[40,55,56,58]. However, Saegusa et. al., demonstrated *α* cell-specific localization of SYT5 in murine islets, whereas Iezzi, et. al, detected SYT5 in both *α* and β cells in rat islets and in the rat INS-1E β cell line. In contrast to both authors, we found that SYT5 showed β cell-specific localization in both murine and rat islets and was absent in *α* cells altogether; overall, the cell-specific expression of SYT5 remains inconclusive. Thus far, our data show a negative relationship between CBD exposure and the transcription of genes important for peptide hormone secretion and Ca^2+^-stimulated cellular excitability, suggesting that CBD may act on multiple transcriptional pathways that intersect in regulating glucagon trafficking and secretion.

One such mechanism we explored was the lysosomal trafficking of glucagon in lysosomes. The retrograde transport of lysosomes is typically associated with lysosomal acidification and degradation of cargo, while anterograde transport is associated with reduced lysosomal acidity, in preparation for membrane repair, or exocytosis[59–61]. In this context, CBD-mediated redistribution of peripheral glucagon-containing lysosomes into the intracellular region suggests that CBD negatively regulates glucagon trafficking, and potentially secretion, by reducing the peripheral density of glucagon available for secretion. These results mirror past work that demonstrated the paracrine factors insulin/GABA and somatostatin rerouted peripheral glucagon intracellularly via the retrograde transport of glucagon in LAMP1^+^ lysosomes[24] in a Stmn2-dependent manner. However, CBD exposure reduced *Stmn2* transcript expression, suggesting that a STMN2-independent mechanism may underlie the CBD-induced redistribution of glucagon.

A potential explanation may lie in functional differences between ECS receptors and those involved in the paracrine regulation of *α* cells. We have previously shown that *Stmn2* knockdown in *α*TC1-6 cells expressing LAMP1-RFP results in peripheral lysosomal accumulation, and that this localization pattern is not further altered by insulin/GABA treatment [24]. In contrast, somatostatin induces the intracellular redistribution of LAMP1-RFP despite STMN2 knockdown, suggesting that somatostatin-induced lysosomal trafficking may function independently of STMN2 [24]. Somatostatin receptors are GPCRs, and GPCRs are known to interact with microtubule machinery to facilitate cargo transport[62]. However, given that the two main canonical ECS receptors are GPCRs, and that CBD has weak activity at both CN1R and CN2R, alternative mechanisms might regulate these changes. Another candidate is the orphaned ECS receptor, known as GPR55. Like the SSTR, GPR55 is a GPCR; however, it is coupled to the G*α*_q_ subunit, which is known to be stimulatory and mediates intracellular Ca^2+^ release from the ER via the conversion of PIP_2_ into IP_3_ by PLC. CBD is a known antagonist of this receptor[49], suggesting that CBD may act on similar downstream pathways that converge on microtubule-associated cargo transport, albeit independently of STMN2 and potentially linked to intracellular Ca^2+^ oscillations; nevertheless, we did not conduct mechanistic studies investigating how ECS-specific agonists and antagonists may affect glucagon trafficking and secretion in the presence of CBD; therefore, the involvement of GPR55 remains speculative. However, this stands as a logical next step in fully elucidating a top-down mechanism linking the ECS, CBD exposure, and glucagon trafficking and secretion.

Current literature is scarce in its characterization of CBD on secretion from neuroendocrine cell types, of which β and *α* cells are categorized. One study, however, investigated insulin secretion in the context of GPR55 antagonism, investigating the potency of CBD to reduce GPR55-stimulated insulin secretion. Using BRIN-BD11 cells, a rat-derived insulin-secreting pancreatic β cell line, researchers demonstrated that GPR55 agonists stimulated insulin secretion in a concentration-dependent, glucose-sensitive manner[7]. CBD, being a GPR55 antagonist, attenuated this response: under high-glucose conditions (16.7 mmol/L), 10 µM CBD reduced abnormal CBD (Abn-CBD)-induced insulin secretion by 56%[7]. CBD alone, at 10 mM, also significantly reduced insulin secretion[7]. These experiments were repeated in isolated islets where, under higher-glucose conditions (11.1 mmol/L), 10 µM CBD reduced Abn-CBD-induced insulin secretion by 12-45%; however, CBD alone did not reduce insulin secretion, suggesting paracrine factor effects on CBD signalling. These findings align with our own, where we saw a significant reduction in glucose-stimulated glucagon secretion from CBD alone under high glucose conditions. Together, these findings suggest that CBD has a suppressive effect on glucose-stimulated insulin and glucagon secretion, and that GPR55 antagonism may be a potential mechanism through which CBD exerts its effects in β cells, and potentially in *α* cells.

While our body of work has provided a comprehensive transcriptomic overview, with observational immunofluorescent microscopy data, and functional secretion data suggesting CBD may act as a negative regulator of glucagon trafficking and secretion, there are a few limitations to the current study. First, exposure to CBD alone in vitro does not fully recapitulate human cannabis consumption in vivo. The ECS in pancreatic islets signals through the release of endogenously produced endocannabinoids from *α*, β, and ∂ cells, which can be altered by the ingestion of cannabis, its active components, Δ9-THC and CBD, as well as their active and inactive metabolites[63]. These components also act on different ECS receptors with different affinities, adding additional layers of regulatory complexity[64–66]. For these reasons, future studies should (i) be conducted in vivo, or in isolated/stem-cell derived pseudo-islets to recapitulate the paracrine actions of the pancreatic islets and of the ECS, (ii) include ECS receptor agonists/antagonists to elucidate receptor-specific effects, and (iii) treat cells with mixtures of Δ9-THC and CBD, mimicking cannabis plant consumption. Nonetheless, we have been able to recapitulate past findings from our *α*TC1-6 cells using in vivo mouse and rat models[22,67,68], supporting our use of the *α*TC1-6 cell line as a model to explore mechanistic outcomes observed in our in vivo rodent models. Additionally, our study included the use of 7-COOH-CBD, an inactive metabolite of CBD, as a secondary control to determine whether CBD-induced changes in glucagon trafficking were due to the pharmacological activity of CBD and rather than non-specific exposure to a structurally related cannabinoid molecule.

In conclusion, our data supports a conceptual framework wherein acute CBD exposure functions as a negative regulator of glucagon trafficking and secretion through the rerouting of peripheral glucagon into lysosome-associated intracellular compartments. We propose that CBD may act through ECS-linked signalling pathways that converge on the regulation of cellular excitability and peptide hormone trafficking, impairing proper *α* cell function, without altering ECS receptor expression or overall ECS composition.

## Supplementary Figures and Tables

**Supplemental Table 1:**
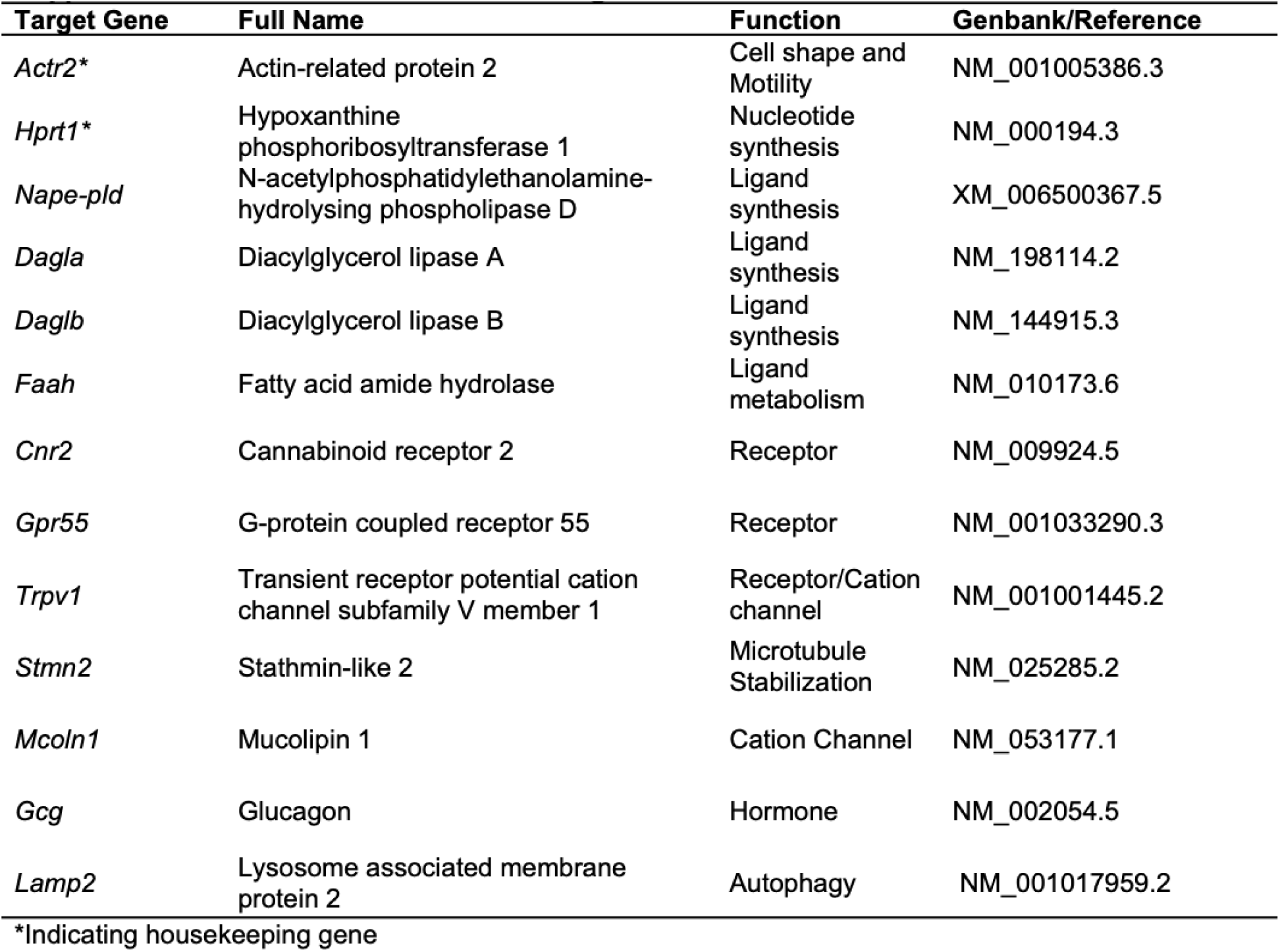
Mouse PCR Primer Targets.

| Target Gene | Full Name | Function | Genbank/Reference |
|---|---|---|---|
| *Actr2\** | Actin-related protein 2 | Cell shape and Motility | NM_001005386.3 |
| *Hprt1\** | Hypoxanthine phosphoribosyltransferase 1 | Nucleotide synthesis | NM_000194.3 |
| *Nape-pld* | N-acetylphosphatidylethanolamine-hydrolysing phospholipase D | Ligand synthesis | XM_006500367.5 |
| *Dagla* | Diacylglycerol lipase A | Ligand synthesis | NM_198114.2 |
| *Daglb* | Diacylglycerol lipase B | Ligand synthesis | NM_144915.3 |
| *Faah* | Fatty acid amide hydrolase | Ligand metabolism | NM_010173.6 |
| *Cnr2* | Cannabinoid receptor 2 | Receptor | NM_009924.5 |
| *Gpr55* | G-protein coupled receptor 55 | Receptor | NM_001033290.3 |
| *Trpv1* | Transient receptor potential cation channel subfamily V member 1 | Receptor/Cation channel | NM_001001445.2 |
| *Stmn2* | Stathmin-like 2 | Microtubule Stabilization | NM_025285.2 |
| *Mcoln1* | Mucolipin 1 | Cation Channel | NM_053177.1 |
| *Gcg* | Glucagon | Hormone | NM_002054.5 |
| *Lamp2* | Lysosome associated membrane protein 2 | Autophagy | NM_001017959.2 |

\*Indicating housekeeping gene

**Supplemental Table 2:**
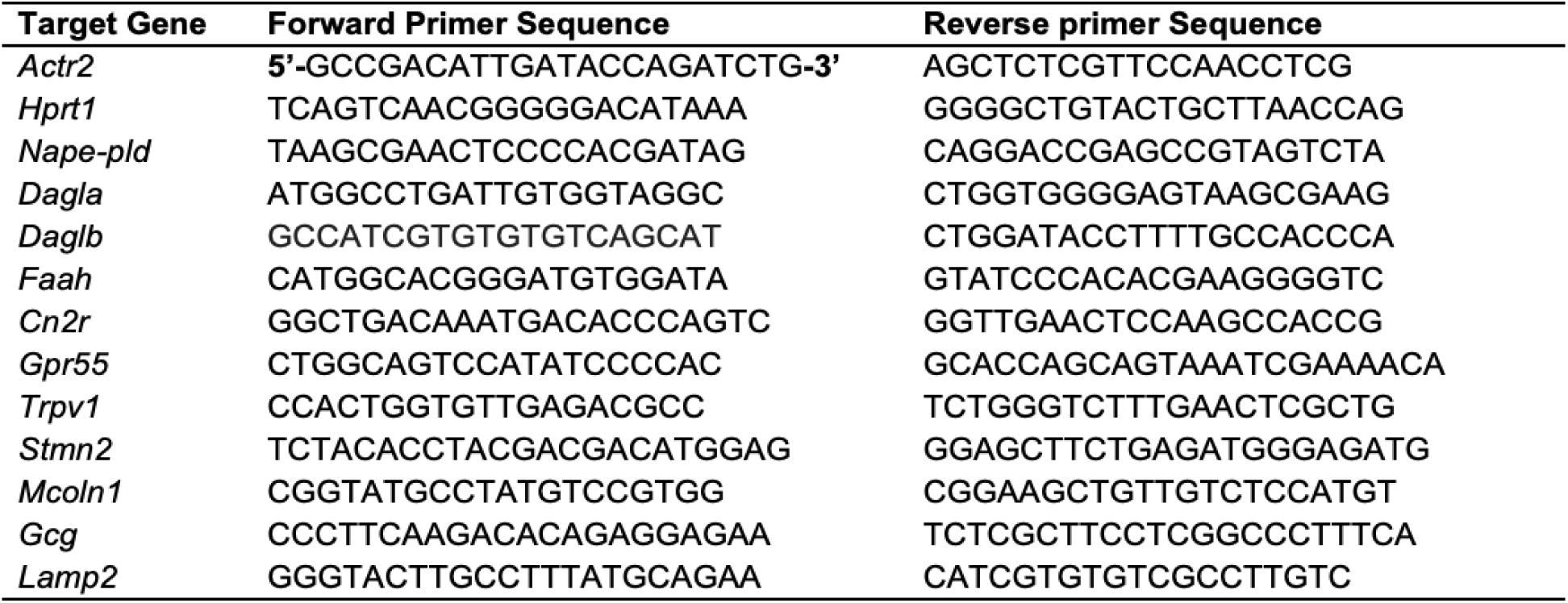
PCR Primer Target Sequences.

**Supplemental Table 3:** Primary and Secondary Antibody Information.

| Antibody | Cell Component | Dilution (V/V) | Manufacturer | Details |
|---|---|---|---|---|
| Mouse anti-glucagon | Glucagon | 1:1000 | Abcam | ab10988 |
| Rat anti-lamp1 | Lysosomal marker | 1:50 | Abcam | ab25245 |
| Rabbit anti-synaptotagminV | Synaptic vesicle marker | 1:500 | Invitrogen™ | PA5-44987 |
| Rabbit anti-syntaxin1a | Plasma membrane | 1:50 | Abcam | ab272736 |
| Donkey anti-mouse IgG Alexa Fluor™ (647) | N/A | 1:500 | Invitrogen™ | A-31571 |
| Donkey anti-rat IgG Alexa Fluor™ (594) | N/A | 1:500 | Invitrogen™ | A-21207 |
| Donkey anti-rabbit IgG Alexa Fluor™ (488) | N/A | 1:500 | Invitrogen™ | A-21206 |

\*Parentheses indicating fluorophore wavelength.

**Supplementary Figure 1:**
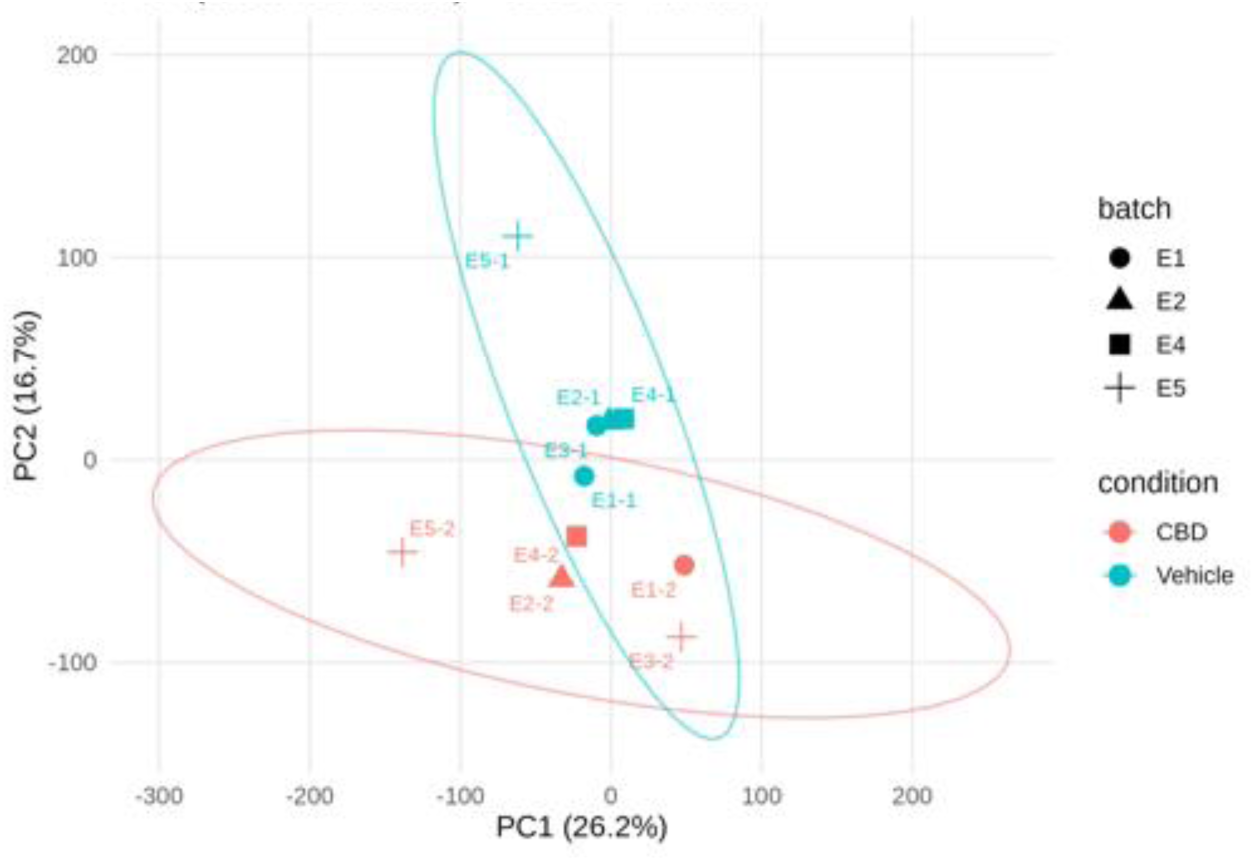
Principal Components Analysis (PCA) Plot between Control and CBD-exposed Groups of *α*-TC1-6 Cells. E1-5 corresponds to the week (batch) the cells were grown.

**Supplemental Figure 2:**
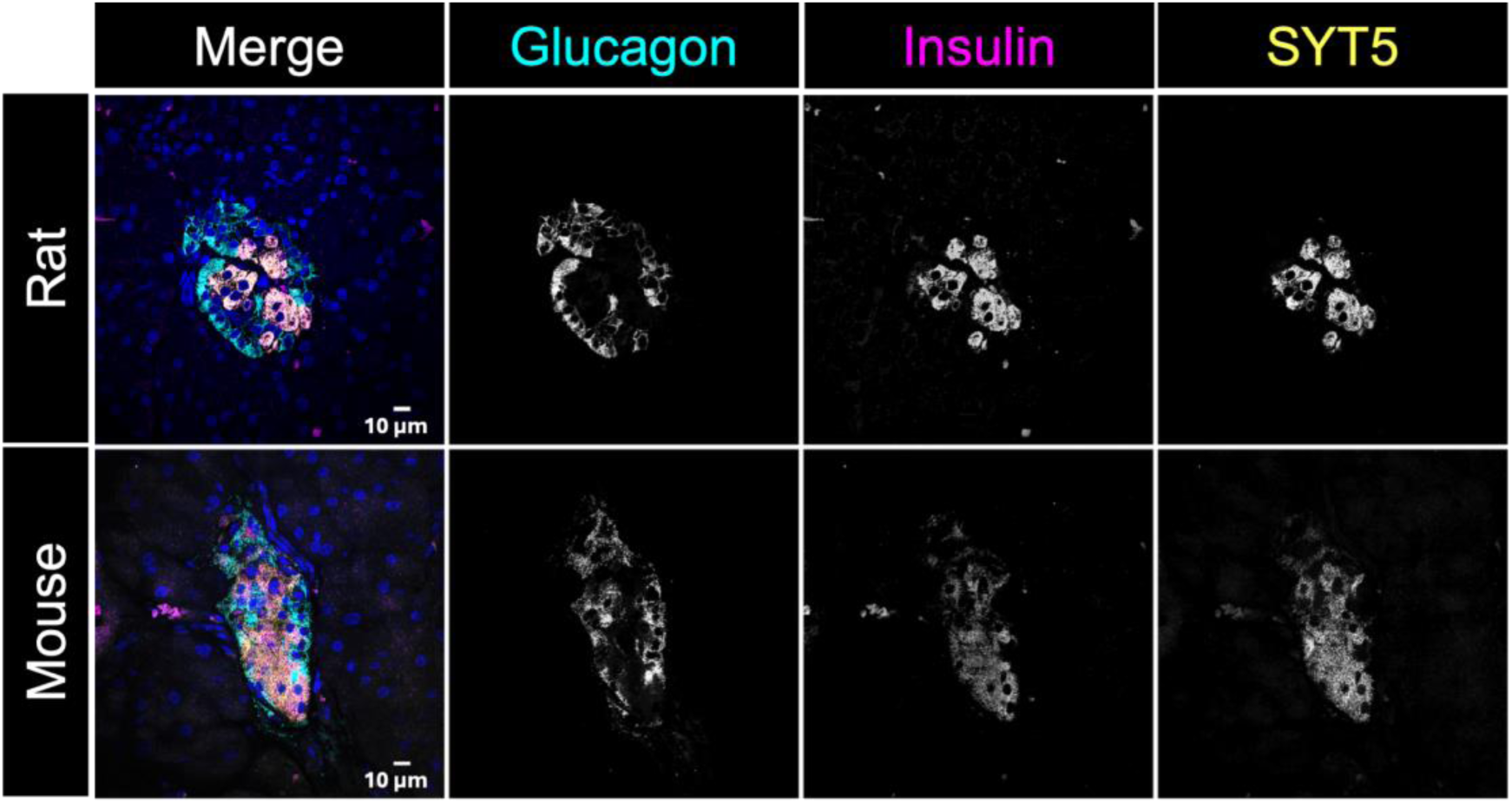
Representative confocal images of SYT5 staining.

**Supplemental Table 4:** M1 MOV Values of Vehicle, CBD, and 7-COOH-CBD Treated *α*-TC1-6 Cells.

|  |  |  |  | Adjusted p-values |  |  |
|---|---|---|---|---|---|---|
| **Comparison** | **Vehicle** | **7-COOH-CBD** | **CBD** | **Vehicle vs. 7-COOH-CBD** | **7-COOH-CBD vs. CBD** | **Vehicle vs. CBD** |
| Glucagon over Lamp1 | 0.361 | 0.344 | 0.450 | 0.901 | 0.004* | 0.015* |
| Glucagon over Syntaxin1a | 0.338 | 0.354 | 0.328 | 0.950 | 0.752 | 0.962 |
| Lamp1 over Syntaxin1a | 0.421 | 0.437 | 0.575 | 0.936 | 0.021* | 0.011* |
| Lamp1 over Syntaxin1a | 0.342 | 0.346 | 0.319 | 0.999 | 0.587 | 0.895 |

\* = Significantly different.

**Supplemental Table 5:** M2 MOV Values of Vehicle, CBD, and 7-COOH-CBD Treated α-TC1-6 Cells.

|  |  |  |  | **Adjusted p-values** |  |  |
|---|---|---|---|---|---|---|
| **Comparison** | **Vehicle** | **7-COOH-CBD** | **CBD** | **Vehicle vs. 7-COOH-CBD** | **7-COOH-CBD vs. CBD** | **Vehicle vs. CBD** |
| Lamp1 over Glucagon | 0.420 | 0.371 | 0.446 | 0.628 | 0.297 | 0.922 |
| Syntaxin1a over Glucagon | 0.356 | 0.406 | 0.371 | 0.486 | 0.447 | 0.975 |
| SynaptotagminV over Lamp1 | 0.242 | 0.241 | 0.310 | 0.999 | 0.067 | 0.076 |
| Syntaxin1a over Lamp1 | 0.258 | 0.274 | 0.330 | 0.978 | 0.573 | 0.445 |

\* = Significantly different.

